# 3D live imaging and phenotyping of the subcellular cytotoxicity in cancer immunotherapy using event-triggered Bessel oblique plane microscopy

**DOI:** 10.1101/2024.03.23.586380

**Authors:** Zhaofei Wang, Jie Wang, Yuxuan Zhao, Jin Jin, Wentian Si, Longbiao Chen, Man Zhang, Yao Zhou, Shiqi Mao, Yicheng Zhang, Chunhong Zheng, Liting Chen, Peng Fei

## Abstract

Clarification of the cytotoxic function of T cells is crucial for understanding human immune responses and immunotherapy procedures. Here, we report an event-triggered Bessel oblique plane microscopy (EBOPM) platform capable of smart 3D live imaging and phenotyping of chimeric antigen receptor (CAR)-modified T-cell cytotoxicity in cancer immunotherapy; the EBOPM platform has the following characteristics: an isotropic subcellular resolution of 320 nm, large-scale scouting over 400 interacting cell pairs, long-term observation across 5 hours, and quantitative analysis of the Terabyte-scale 3D, multichannel, time-lapse image datasets. Using this advanced microscopy platform, several key subcellular events in CAR-T cells were captured and comprehensively analyzed; these events included the instantaneous formation of immune synapses and the sustained changes in the microtubing morphology. Furthermore, we identified the actin retrograde flow speed, the actin depletion coefficient, the microtubule polarization and the contact area of the CAR-T/target cell conjugates as essential parameters strongly correlated with CAR-T-cell cytotoxic function. Our approach will be useful for establishing criteria for quantifying T-cell function in individual patients for all T-cell-based immunotherapies.

## Main

Immunotherapy has undoubtedly revolutionized cancer treatment^1–4^. For instance, chimeric antigen receptor (CAR)^5^-engineered T cells (CAR-T cells) have ushered in a new era for treating hematological malignancies. However, highly selected patients still have overall response rates (ORRs) ranging from 50-90%^6^. Furthermore, only a small proportion of patients achieve long-term remission from immunotherapy^7^. Visualization and quantitively analysis of the cytotoxic activity of T cells derived from individual patients by fluorescence microscopy could be a potentially effective approach for predicting the efficacy of T-cell-based cellular immunotherapy^8^. However, current functional studies of CAR-T cells have focused mostly on the typical attributes of cell populations and lack long-term phenotyping of antitumor cytotoxicity at the subcellular level; the latter has great potential for the identification of the efficacy-related biomarkers.

In contrast to classical fluorescence microscopy techniques, light-sheet fluorescence microscopy (LSFM) can rapidly image live cells in three dimensions (3D) with low phototoxicity, thereby allowing for a more physiologically relevant assessment of T-cell functionality^8^. The LSFM not only intuitively reveals the dynamic physiological processes of cell toxicity^9–11^ but also facilitates the regression analysis of the cell states through the feature analysis of the images^12^. However, due to the absence of an efficient sample mounting and data analysis pipelines, current LSFM techniques are regarded as low-throughput imaging techniques limited to the observation and analysis of individual cells^12^ and cannot be used to study large-scale cell events at high throughput. Due to the heterogeneity of CAR-T cells^13,14^, the results from the analysis of the individual cells potentially exhibit significant discrepancies in overall performance. Therefore, a high-throughput long-term live-cell imaging system is needed for analyzing the CAR-T-cell cytotoxicity.

Previous studies reported that the quality of the CAR-mediated immunological synapse (IS) was essential for CAR-T-cell effectiveness^15–17^. However, the evaluated ISs were all from fixed CAR-T/target conjugates. The evaluation of IS formation and subsequent dynamic changes in CAR-T cells has never been reported. In this study, we developed a high-throughput, high-resolution, multidimensional live-cell phenotyping pipeline consisting of the following: a dedicated microfluidic chip for large-scale pairing of the CAR-T/tumor cells; an event-triggered Bessel oblique plane microscopy (EBOPM) that could automatically detect immune cytotoxicity events and three-dimensionally image the subcellular interactions at high spatiotemporal resolution over a long tumor killing time of hours; and a comprehensive spatial-spectrum cell phenotyping algorithm that could specifically identify and quantitatively analyze organelle phenotypes that were strongly correlated with the killing efficacy. With this microfluidics-enabled intelligent EBOPM platform, we revealed the dynamics of immune synapses as key factors for evaluating the cytotoxicity of CAR-T cells.

## Results

### Overview of event-triggered Bessel oblique plane microscopy (EBOPM)

We designed an open-environment microchamber chip containing 2000 cylindrical chambers (50-μm diameter × 50-μm depth) to accommodate the CD3+ T cells transduced with concentrated CAR19-Lifeact-EGFP lentivirus and Nalm6 tumor cells with membranes labeled with mApple **(Fig. 1a)**. The CAR-T and Nalm6 cells were mixed 1:1 on the chip to form large CAR-T-Nalm6 cell pairs in the microchambers **(Fig. 1b, Supplementary Fig. 1)**. Notably, we fabricated the microchip using Bio-133 glue, a material with a water-like refractive index of 1.33, to ensure imaging without aberration from the microchambers^19^. We devised a smart imaging algorithm based on automatic image scoring to enable instant positioning of the preferred microchambers, where CAR-T cells were paired with tumor cells one on one **(Fig. 1c, Supplementary Fig. 2)**.

**Figure 1.**
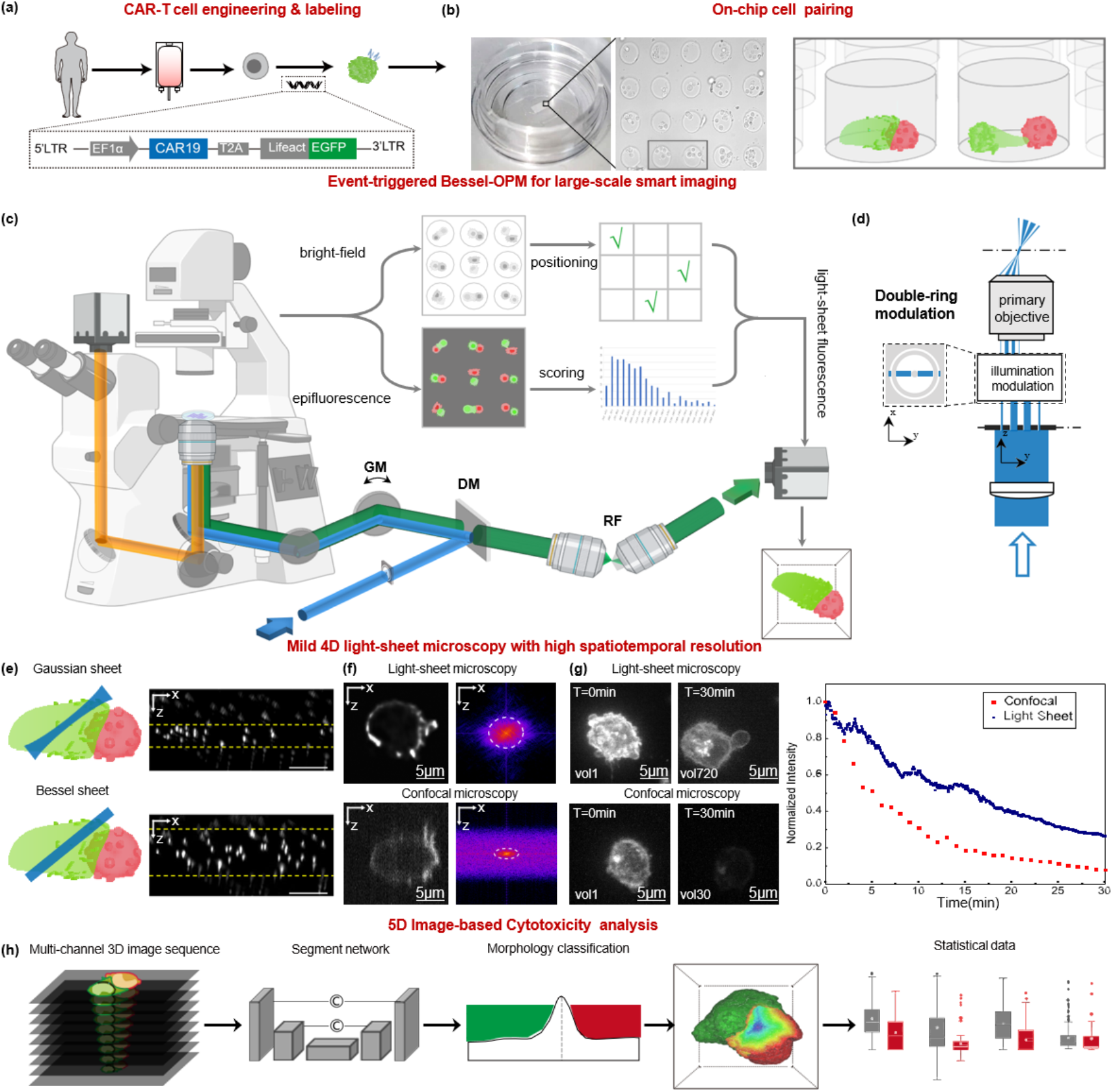
Setup of the event-triggered Bessel oblique plane microscopy (EBOPM) (a) Cell engineering to prepare CAR-T cells transfected with the CAR19 and EGFP genes. (b) Microchamber chip structure diagram. The cells were loaded into the chip. (c) EBOPM composed of smart imaging algorithms, the Bessel-OPM optical system and cytotoxicity analysis algorithms. GM: galvo mirror. DM: dichroic mirror. RF: remote focus system. (d) Double-ring modulation to generate ultrathin Bessel light sheets with tunable axial extents from 0.5 to 1.5 μm. (e) Comparison between Gaussian and Bessel light sheets. (f) Comparison of axial resolution between the confocal microscope and our EBOPM. On the left, CAR-T-cell fluorescence images are shown, and on the right, the corresponding Fourier transforms are shown. (g) Comparison of photobleaching between confocal microscopy and our approach. OPM imaged 720 volumes within 30 minutes, while confocal imaging only imaged 30 volumes. (h) Cell phenotype analysis pipeline containing the image reconstruction, segmentation, identification and analysis.

To three-dimensionally image the large-scale antitumor cytotoxic effects of these cell pairs, we developed a new event-triggered Bessel OPM based on the latest OPM setup^15^ **(Fig. 1c, Supplementary Fig. 3)**. We used advanced double-ring light-sheet modulation^18^ (**Fig. 1d**) to allow on-chip light-sheet excitation/detection from the same objective, providing an extended illumination length and sharp optical sectioning suitable for imaging floating CAR-T cells (**Fig. 1e**). Target cells and CAR-T cells were loaded on one chip in sequence, and this intelligent EBOPM rapidly identified 40 qualified wells and recorded their entire 4D cytotoxic events. We designed an axial-to-lateral isotropic deep learning network (**Supplementary Fig. 4**) to reconstruct the light-sheet image sequence of cell interactions with a 3D isotropic resolution of ∼320 nm (**Fig. 1f**) and low phototoxicity for over 700 measurements; these were both superior to line-confocal microscopy (compared in **Fig. 1g**).

**Figure 2.**
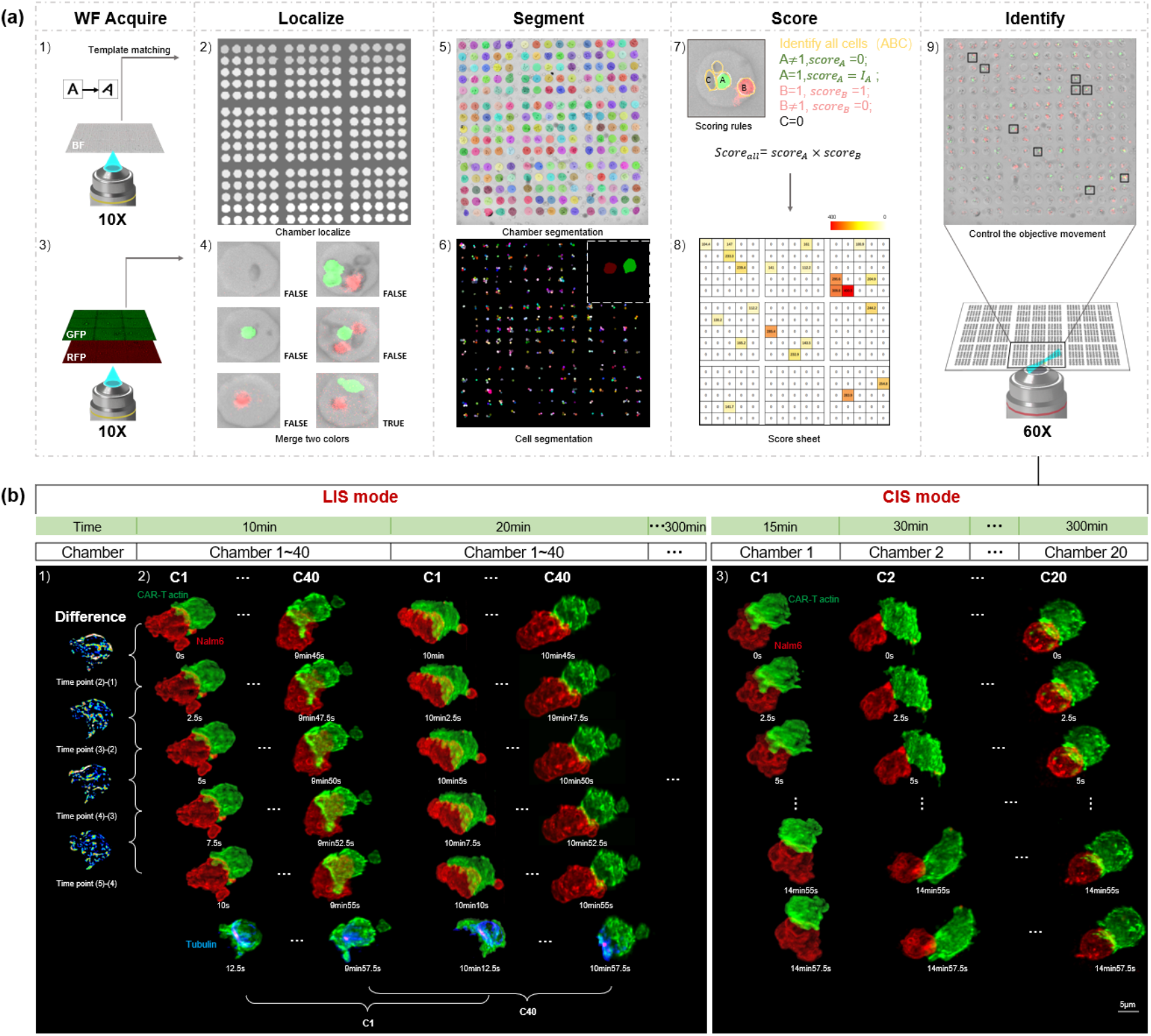
Smart imaging procedure and imaging strategy. (a) Smart imaging procedure. 1) BF mode with a 10× objective. 2) Localization of all microchambers based on wide-field images using the smart imaging algorithms. 3) Wide-field fluorescence mode with a 10× objective. 4) Example of evaluation criteria. “TRUE” indicates a qualified chamber. 5) Chamber segmentation. 6) Segmentation of all cells via deep learning. 7) Scoring criteria. 8) Score table of each chamber. 9) Identification of the chambers with highest scores. The microscope was switched to OPM mode. (b) Continuous and long-term imaging strategy (CIS & LIS). 1) Differences between the adjacent volumes. 2) LIS mode. The algorithm automatically selected 40 microchambers, completed a scan round within 10 minutes, and imaged 6 volumes in 15 s per microchamber. The first five volumes contained three-channel information on CAR-T-cell actin, Nalm6 cell membranes and SYTOX blue, while the last volume contained CAR-T-cell actin and CAR-T-cell tubulin. 3) CIS mode. The algorithm selected several microchambers and continuously imaged each for 15 minutes. All volumes contained three-channel information on CAR-T-cell actin, Nalm6 cell membranes and SYTOX blue.

We also developed a phenotype analysis pipeline for automatically processing these terabyte images with high throughput **(Fig. 1h, Supplementary Figs. 5-7)**. This pipeline could automatically extract the numerical features from the cells for analysis, thereby evaluating the cytotoxic activities through quantifying CAR-T actin’s motion, their contact areas with the tumor cells and the polarity of the microtubule organizing center (MTOC).

### Event-triggered smart imaging algorithms and imaging strategies

In the smart imaging procedure (**Fig. 2a**), we used bright-field mode to quickly obtain images of the chip (**Fig. 2a1**) and then used the template matching algorithm to discern the coordinates of all the microchambers (**Fig. 2a2**). Furthermore, in wide-field fluorescence mode (**Fig. 2a3**), we detected chambers with the desired cell pairs (**Fig. 2a4**) by deep learning segmentation and scoring algorithms (**Fig. 2a5∼9**). We segmented the bright-field fluorescence composite image into small blocks based on the localization results (**Fig. 2a5**) and then used CellPose^20^ deep learning segmentation to identify the cells in each microchamber (**Fig. 2a6**). Based on the above segmentation results, we could quantify the cell distribution in each microchamber. As shown in **Fig. 2a7**, cell A and cell B, which had fluorescence signals, were recognized as CAR-T cells and tumor cells, respectively, while the remaining cells without fluorescence were considered T cells that were not engineered with the CAR gene. When the number of CAR-T cells or tumor cells in the microchamber was not equal to 1, the microchamber score was set to zero. Otherwise, we scored the chamber based on the fluorescence intensity of the CAR-T cells. Thus, we obtained a scoring table for all microchambers (**Fig. 2a8**). Finally, the coordinates of the selected microchambers were returned to the control software for light-sheet imaging (**Fig. 2a9**) in continuous or long-term mode.

To capture both the cellular motion details and long-term features, we applied continuous imaging for 15 minutes and long-term imaging for 5 hours. Using a continuous imaging strategy (CIS, **Fig. 2b1**), we three-dimensionally imaged the interactions of the paired CAR-T cells/target cells in each chamber at a rate of 0.4 volume/s for 15 minutes. Under a long-term imaging strategy (LIS **Fig. 2 b2**), we also captured the images at a rate of 0.4 volume/s for 15 s each round and then switched to the next cell. After 40 chosen cells were imaged, they were returned to the first cell, and the above process was repeated, with each imaging cycle lasting 10 minutes (15 s/cell × 40 cells). LIS not only improved the imaging throughput but also notably reduced the phototoxicity to cells, extending the imaging window of CAR-T cells from a few minutes^21^ to several hours until the endpoint was reached. The combination of CIS and LIS thereby established cross-validation for compressive visualization of the cytotoxic activities.

### Imaging of the CAR-T/Nalm6 immune synapse (IS) formation

The immune synapse is known to play a crucial role in the cytotoxicity of both the T-cells and CAR-T cells^15–17,22^. With intelligent EBOPM imaging, we first performed high-throughput 5D (space + time + spectrum) imaging of early immune synapse (IS) formation in real CAR-T/Nalm6 interactions. Previous studies suggested that when T cells formed immune synapses, they usually contacted the target cells on the dense actin side, followed by the polarized movement of the MTOC toward the target cells^9^; however, based on our results (**Fig. 3a**), the CAR-T cells tended to establish immune synapses on the MTOC side, usually on the sparse actin side. Subsequently, instead of MTOC, actin polarized toward the target cell, aggregated around the target cell to form the IS, and ultimately completed the cytotoxicity process (**Fig. 3b**). Interestingly, when CAR-T cells were treated with dasatinib^23^, a drug reported to induce a function-off state in CAR-T cells, their cytotoxicity appeared to be blocked at the contact state, as shown in **Fig. 3b**. These CAR-T cells tended to contact and adsorb to the tumor cells on the MTOC or sparse actin side, but the actin polarity reversal and an effective IS formation was suppressed. (**Fig. 3c**). Through CIS, we recorded the actual process of actin polarity reversal when the CAR-T cells formed immune synapses (**Fig. 3d**). We observed that 62.8% of the cells followed the pattern shown in **Fig. 3b**, and 28.2% of the cells did not undergo significant polarization before forming immune synapses. Only 9% of the cells formed immune synapses on the dense actin side, after which the MTOC migrated (**Supplementary Fig. 8**). These results provide new insight into CAR-T/target cell IS formation.

**Figure 3.**
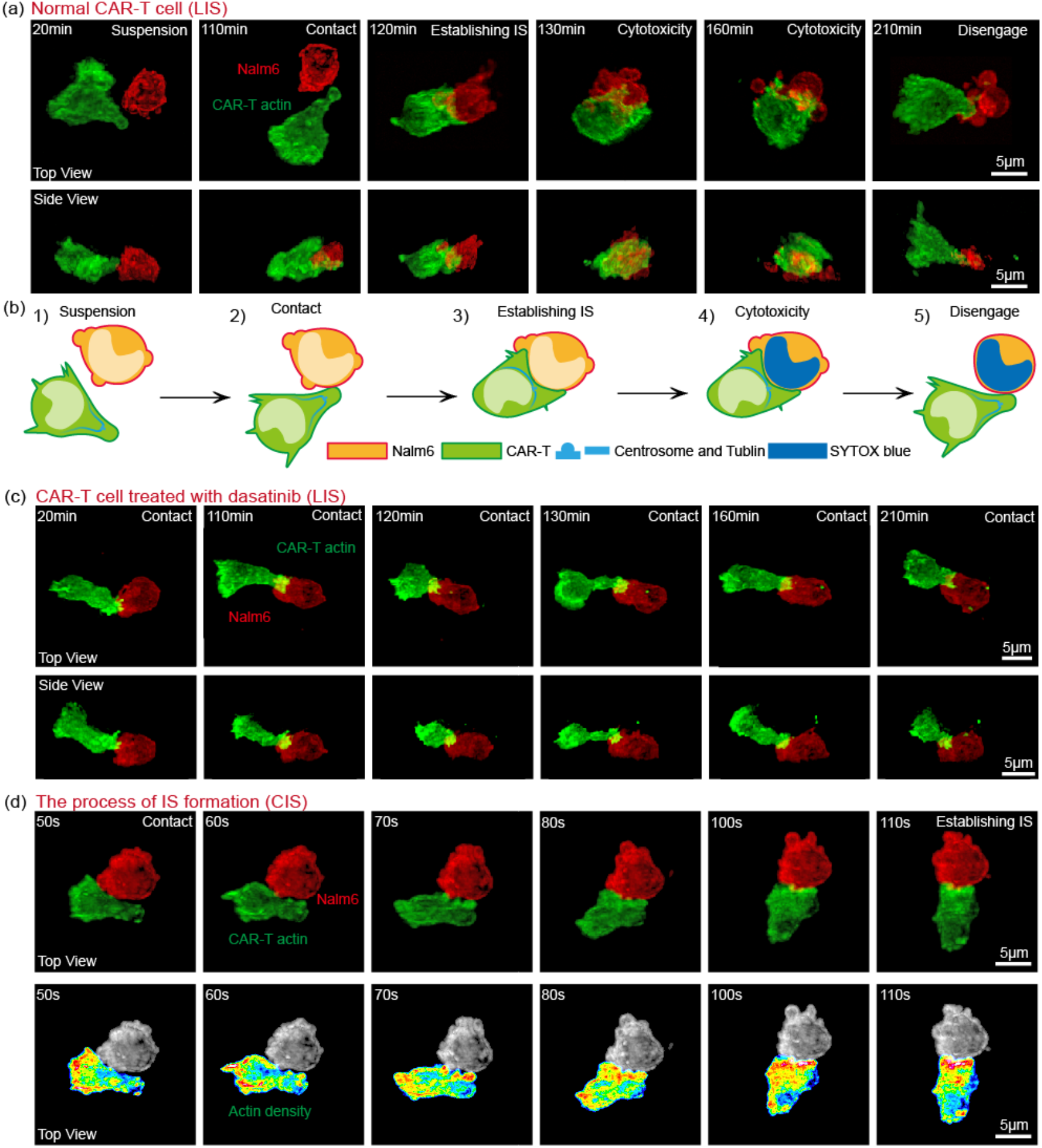
Cytotoxic behavior. (a) A normal CAR-T-cell captured by the EBOPM under a long-term imaging strategy (LIS). The upper column is the top view of the 3D image, and the lower column is the side view. (b) Schematic diagram of a typical cytotoxic behavior pattern. 1) During the separation stage, some CRA-T cells undergo morphological polarization. 2) Polarized CRA-T cells tend to contact the target cells with sparse actin. 3) Actin density polarity undergoes reversal, and IS is established. 4) Cytotoxic process killing target cells. 5) CAR-T cell polarity restoration and attempt to disengage from target cells. (c) CAR-T cells treated with dasatinib captured by EBOPM under LIS. Its cytotoxic behavior was blocked at the contact stage. (d) Normal CAR-T-cell captured by the EBOPM under a continuous imaging strategy (CIS). The above column displays the original fluorescence image, while the following column uses a heatmap to illustrate the change in the polarity of the actin density.

### Cytotoxicity analysis algorithms (CAAs)

To analyze a large amount of 5D image data, we developed a CAAs for the automatic quantification of the cell features **(Fig. 4)**. The raw light-sheet imaging data were initially subjected to the reconstruction algorithm (RA) to restore the spatial relationship of the OPM **(Fig. 4a, b)**; then, the 5D image data were subjected to a quantitative algorithm (QA) to obtain the quantitative indicators of the cells **(Fig. 4c, d)**. Based on previous reports, we selected 5 QA indicators to quantify the characteristics of the CAR-T cells, including the area of the IS, actin retrograde flow speed, actin depletion coefficient of the IS, MTOC polarization angle, and tumor cell death rate. The tumor cell death rates could be directly determined based on SYTOX BLUE staining **(Fig. 4c)**, while the remaining indicators required algorithms to extract the information from the images.

**Figure 4.**
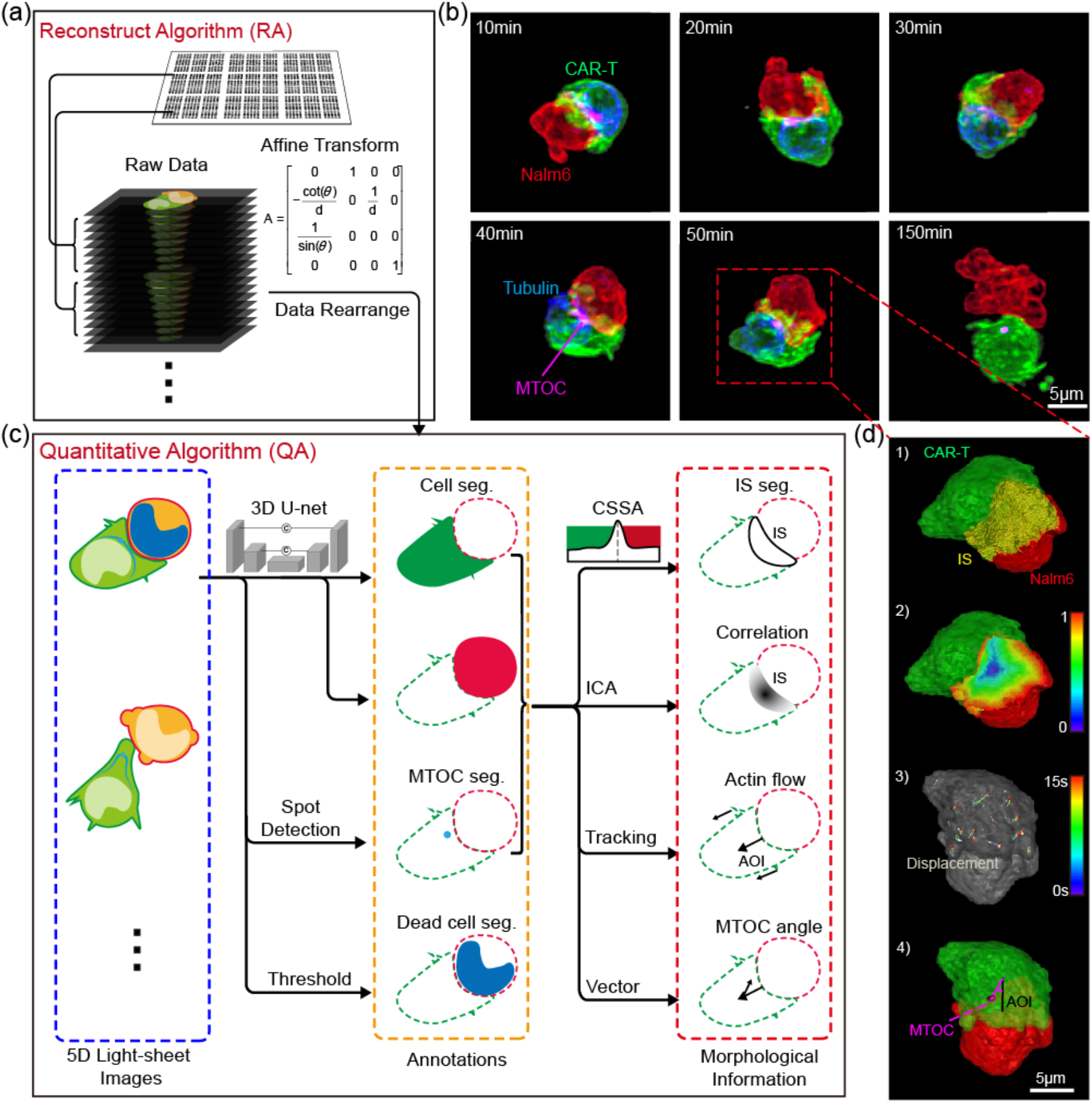
Cytotoxicity analysis algorithms (CAAs). (a) Reconstruction algorithm (RA). The raw image data captured by the EBOPM required data rearrangement and affine transform batch processing to calibrate the tilted spatial relationship of the EBOPM. (b) Reconstructed 5D image data (space + time + spectrum). (c) Quantitative algorithm (QA). CSSA: contact surface segmentation algorithm. ICA: iterative corrosion algorithms. (d) Visualization of intermediate results in QA. 1) The segmentation of the immune synapses by CSSA. 2) Normalized edge distance of the immune synapses via the ICA. 3) Actin flow trace via a tracking algorithm. 4) Visualization of the MTOC polarization angle.

Prior research on IS has focused primarily on the 2D IS formed by the T cells on the surface of antigen-coated slides ^15,24,25^. The QA introduces a contact surface segmentation algorithm (CSSA), which can accurately segment the 3D contact surface with a thickness of 1 pixel, and the area of the interactive IS can then be obtained **(Fig. 4d 1, methods, Supplementary Fig. 6)**. Furthermore, we developed an iterative corrosion algorithm (ICA) to obtain the distance from every segmented pixel on the IS to its center and calculated the actin depletion coefficient *r* between this distance and the corresponding fluorescence signal intensity of F-actin **(Fig. 3d 2, methods, Supplementary Fig. 6)**.

In both cytotoxic T lymphocytes (CTLs) and natural killer (NK) cells, robust actin polymerization-driven retrograde actin flow at the perimeter of the IS was observed^26,27^. However, whether the actin retrograde flow occurred in CAR-T cells and its role in the effectiveness of CAR-T cells were not clear^9,12^. In QA, we defined the vector pointing from the center of the IS to the center of the CAR-T-cell as the axis of interaction (AOI), identified and tracked the dendritic actin structures on the T-cell surface, recorded the displacement vectors **(Fig. 4d 3)** of these structures within the 15 s imaging window, and statistically measured the intensity of the projection of these displacement vectors onto the AOI vector to extract the eigenvalues of actin flow during this period. We also calculated the angle between the AOI and the vector from the MTOC to the center of the CAR-T cells **(Fig. 4d 4)**, which was termed the MTOC polarization angle, to investigate the reorientation of the MTOC during the cytotoxic process^28^.

### Quantitative analysis using the CAAs

**Fig. 5a** shows the actin retrograde flow speed of the CAR-T cells from **Fig. 3a**. A correlation was observed between the flow speed and the cytotoxicity stage. **Fig. 5b** shows the visualization of the actin movement traces at the peak flow speed in the state of established IS; these results indicated that actin tended to flow in the opposite direction of the immune synapses.

**Figure 5.**
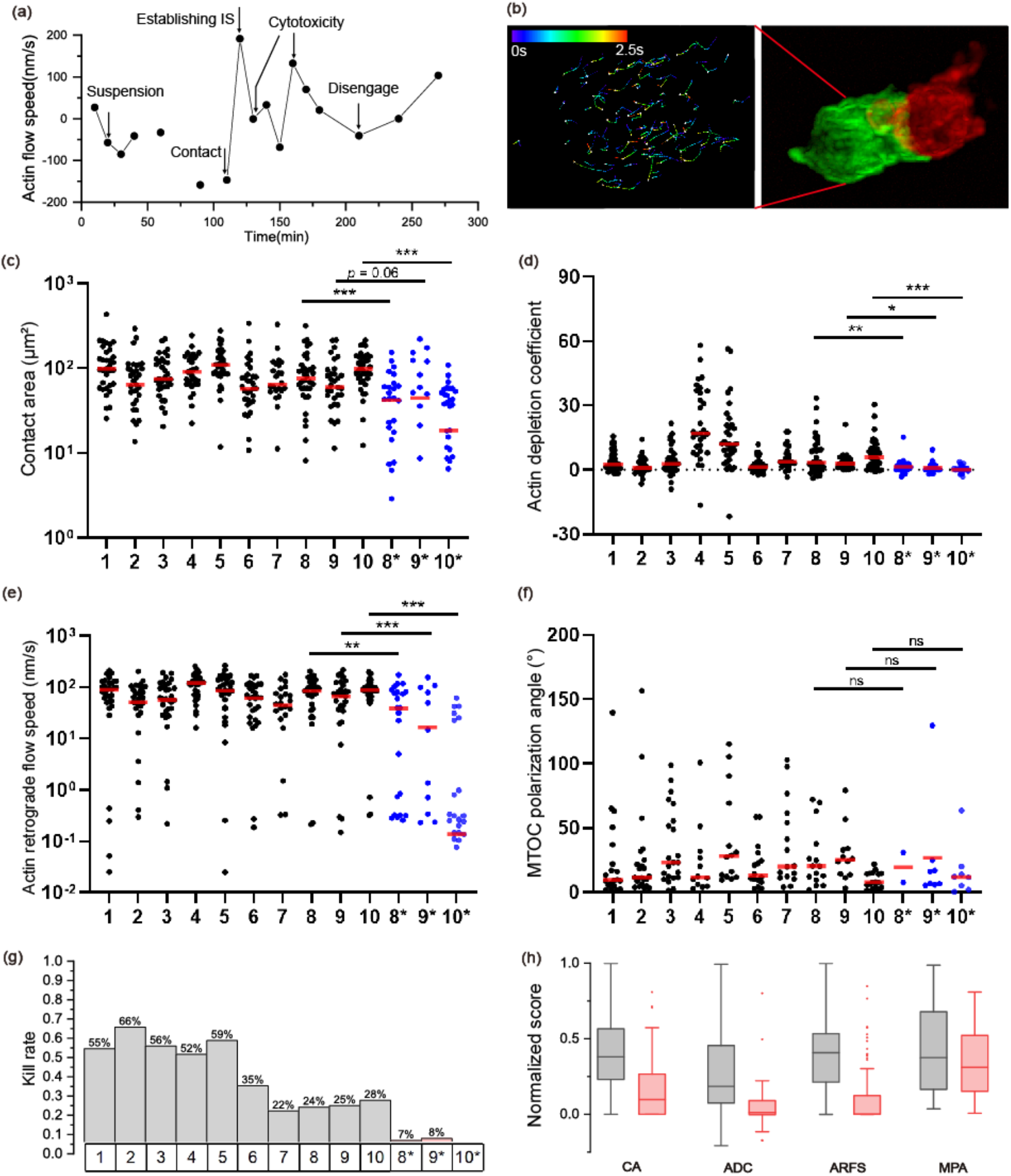
Cytotoxicity quantitative statistical analysis. (a) Use of the algorithm to quantify the actin flow velocity shown in Fig. 3 a. (b) Actin flow traces at 120 minutes. (c) Graph of the IS area. (d) Graph of the actin depletion coefficient between the normalized edge distance and actin fluorescence intensity. (e) Graph of actin flow speed. (f) Graph of the MTOC bias angle. (g) Graph of the mortality rate of Nalm6 cells labeled with SYTOX blue. (h) Summary graphs of the data from Fig. c∼f, where the cells treated with dasatinib are red and the control cells are gray. ^*^: dasatinib-treated group.

Using the CAAs, we analyzed all data. We imaged CAR-T cells from 10 healthy donors to determine the baseline. Three of the ten donor-derived CAR-T cells were also imaged after treatment with dasatinib to test the sensitivity of specific indicators to CAR-T-cell function. **Fig. 5c-e** shows the statistical results from the area of IS, the actin depletion coefficient of IS and the actin retrograde flow speed, and all three statistical indicators showed a significant decrease in dasatinib-treated cells. The MTOC polarization angle showed a distribution trend of less than 90 degrees, but no significant difference was observed in this indicator between the dasatinib-treated CAR-T cells and the controls **(Fig. 5f, Supplementary Fig. 9**); these results indicated that dasatinib might not block the physiological process of MTOC polarization. Moreover, the tumor cell death rates were also significantly reduced in the dasatinib-treated cells (**Fig. 5g**). We merged data from different sources of healthy donors to obtain a distribution box chart for the four indicators mentioned above, with the exception of the tumor cell death rates, to visually demonstrate the above conclusions.

## Discussion

In summary, based on the new development of intelligent EBOPM and T-cell behavior analysis, we accomplished high-throughput, long-term 3D imaging and phenotyping of CAR-T/Nalm6 interactions to comprehensively analyze the antitumor cytotoxic function of CAR-T cells. The advanced imaging technique in conjunction with high-dimensional image-based analysis formed an automated and scalable cell observation and analysis pipeline well suited for diverse types of cell research. In our study, we tested the ability of this system to evaluate the function of CAR-T cells in specific cases. The visualization and statistical results were significantly different between the dasatinib-treated CAR-T cells and controls, validating the effectiveness of our platform.

Our method has broad research prospects. From an algorithmic perspective, extracting KB-level information from TB-level data not only the five indicators proposed in this study. For example, in our result, we found that there may be errors in using dyes (SYTOX™ Blue or PI) to indicate whether target cells were killed. The apoptosis of target cells often starts with the morphological changes. Some cells expand and become round, some cells break into pieces, and dead and live dyes usually need to be delayed for several hours to cause the emission of fluorescence from the dead target cells. In our study, the death time of the target cells was not calculated, but the death time of the target cells determined from a morphological perspective may have research value. In addition, many other phenomena, such as microtubule morphology distribution and the CAR-T-cell length-to-diameter ratio, could be statistically analyzed. From a cellular physiological perspective, why do CAR-T cells tend to form immune synapses on the centrosome side? Is this related to receptor distribution? Our study focused on the methods, and we have not yet conducted in-depth research on the abovementioned aspects.

Our method still has limitations. Thirty minutes is needed for the cells to mix and begin imaging. The capturing of the early events of the immune synapse formation on a larger scale still needs further optimization. In addition, the optimization of the analysis algorithm is insufficient, and there is still room for improvement in the 8-hour running time.

Notably, all killer cell-based immunotherapies should have the potential to use our platform for an in-depth study of the microscopic behaviors of cells. Since the IS contact area and actin flow speed were identified as the key markers strongly related to the killing efficacy of CAR-T cells, we anticipate that our approach can effectively advance cancer immunotherapy and support clinical trials in the future.

## Acknowledgements

The authors would like to thank Dr. Meng Zhang, Prof. Yuhui Zhang, and Prof. Liang Huang for providing us insightful biomedical supports, thank Optoelectronic Micro&Nano Fabrication and Characterizing Facility of Huazhong University of Science and Technology for providing technical support on photolithography technology. This work was supported by the funding from National Natural Science Foundation of China (T2225014, 82270238, 21927802). National Key Research and Development Program of China (2022YFC3401102).

## Author contributions

P.F., L.C. and Z.W. conceived the idea. P.F. and L.C. oversaw the project. J.J., M.Z. and S.M. conducted the cell experiments. Y.Z., J.W., Y.Z. developed the optical setups and acquired the experimental images. Z.W., W.S., L.C. developed the programs and processed the images. P.F., L.C., Z.W., W.S. and J.J. analyzed the data. Z.W., J.W., L.C., Y.Z., C.Z. and P.F discussed and wrote the paper.

## Methods

### Cell lines and cell culture

The acute B-lymphocytic leukemia cell line Nalm6 was cultured in RPMI 1640 medium (Gibco, Grand Island, NY, USA) supplemented with 10% fetal bovine serum (FBS; Gibco, Grand Island, NY, USA). The lentivirus packaging cell line LentiX™293T was cultured in Dulbecco’s modified Eagle medium (DMEM) (Gibco, Grand Island, NY, USA) supplemented with 10% FBS. The strains were obtained from the American Type Culture Collection (ATCC) and authenticated by short tandem repeat (STR) analysis before use.

### Recombinant plasmid construction and lentivirus packaging

The anti-CD19 scFv (single-chain variable fragment) was grafted into a second-generation CAR with a CD8a hinge/transmembrane region, a CD28 costimulatory domain, and an intracellular CD3ζ. The scFvs were derived from the FMC63 clone under patent WO2012079000^29^. Then, the CD19 CAR gene was linked to Lifeact-EGFP^30^ via the T2A sequence to facilitate *in vitro* visualization. mTagRFP-Membrane-1 was a gift from Michael Davidson (Addgene plasmid # 57992), and we replaced mTagRFP with mApple to construct the mApple-Mem plasmid. Recombinant plasmids were extracted by using the EndoFree Plasmid Maxi Kit (Qiagen, Hilden, Germany) according to the manufacturer’s instructions. Lenti-X™293T cells were co-transfected with a vector carrying the target gene (CD19 CAR-Lifeact-EGFP or mApple-Mem) and the psPAX2 and PMD2.G packaging plasmids using the transfection agent Lipofectamine 3000 (Invitrogen, Waltham, MA, USA). The viral supernatants were collected, filtered, and concentrated 48 hours after transfection by ultracentrifugation (Avanti J-26S XPI) and then aliquoted and stored at -80 °C.

### Isolation, activation, transduction and culture of T cells

Peripheral blood samples were taken from 10 healthy donors (HDs). Peripheral blood mononuclear cells (PBMCs) were isolated by density gradient centrifugation on Ficoll-Paque Plus (GE Healthcare, Boston, MA, USA). CD3+ T cells from PBMCs were separated using CD3 microbeads (Miltenyi Biotec, Bergisch Gladbach, Germany) following the manufacturer’s instructions. Then, the T cells were stimulated with Dynabeads™ Human T-Activator CD3/CD28 (Gibco, Grand Island, NY, USA) at a 1:1 ratio in CTS™ OpTmizer™ medium (Gibco, Grand Island, NY, USA) containing 5% human AB serum, 2 mM l-glutamine (Gibco, Grand Island, NY, USA) and 200 IU/mL rhIL-2 (PeproTech, Rocky Hill, NJ, USA). Within 24 hours, primary T cells were transduced with concentrated lentivirus at a certain multiplicity of infection (MOI) ranging from 2 to 5. Twenty-four hours later, the T cells were centrifuged and resuspended in fresh culture medium at a density of 1-2×10^6^/mL. Imaging and functional assays were performed after 12 days of culture *in vitro*.

### Construction of mApple-Mem-expressing Nalm6 cell lines

Nalm6 cells were transduced with concentrated mApple-Mem lentivirus at a certain MOI in the range of 30 to 50 for 4 hours. mApple-positive cells were sorted 5 days after infection on a Moflo XDP flow cytometer (Beckman Coulter) to obtain mApple-Mem-expressing Nalm6 cell lines.

### Cell preparation for imaging

For experiments on the impacts of dasatinib, CAR-T cells were pretreated with 50 nM dasatinib (Selleck, Shanghai, China) for 24 hours. Due to the reversible effect of dasatinib, 50 nM dasatinib was also added to all subsequent staining, imaging, and other experimental solutions.

To label the microtubules, CAR-T cells were stained with a SiR-tubulin probe (SpiroChrome, Switzerland) to a final concentration of 2 μM and incubated for 1 hour in a humidified 5% CO_2_ incubator at 37 °C^31^. The cells were then washed twice with warm phosphate buffer saline (PBS) and resuspended in an imaging solution consisting of phenol red-free 1640 medium (Gibco, Grand Island, NY, USA) supplemented with 10% FBS, 25 mM N-2-hydroxy-ethylpiperazine-N′-2-ethanesulfonic acid (HEPES, Gibco, Grand Island, NY, USA), 100 U/mL penicillin and streptomycin (Gibco, Grand Island, NY, USA), and 1 μM SYTOX™ Blue stain (Invitrogen, Waltham, MA, USA).

### Microchip preparation

The first step was to make the mold plate. We designed and customized a polyester mask (SI Fig. 2). Then, we used a photoresist (SU8-2050, Kayaku Advanced Materials) to spin-coat a 50 µm thin layer on a 2-inch wafer and used a UV exposure machine (SUSS MicroTec MA/BA8) to transfer the pattern on the polyester mask to the photoresist; this was developed and used as the mold plate.

The second step was to reverse the mold. The UV-curable adhesive (Bio-133, MyPolymers) was added dropwise onto the mold plate, and the top was flattened as much as possible using a coverslip. After sufficient UV exposure, the cured chip was carefully removed, and the demolded chip was immersed in anhydrous ethanol for more than 24 hours to remove the residual adhesive^32^. The chip was placed at the bottom of a confocal Petri dish and pressed to fix it. Before use, the chip was immersed in sterile water for more than 12 hours to expel the gas inside the chamber, and the use of a vacuum pump to create a low-pressure environment could accelerate the process.

T-cell medium was used to replace the sterile water, 500 μL of the medium was added to the confocal dish to submerge the chip, and then 60 μL of CAR-T cells was added dropwise at a density of 1×10^6^/mL. The plate was incubated for 10 minutes to allow the cells to fall into the chamber. Subsequently, an equal amount of target cells was removed, and the above procedure was repeated. Theoretically, the distribution of cells satisfies the binary Poisson distribution, and λ needed to be set to approximately 1. In actual experiments, the smart imaging algorithms estimated λ, which was convenient for the adjustment of the cell density in real time.

### Optical system

We built a single-objective light sheet imaging system using an Olympus ix83 microscope. The system magnification was 63.3×, and the primary objective (O_1_) used a 60×1.3NA silicone oil objective (UPLSAPO60XS2, Olympus). O_1_ was combined with a 40×0.95NA air objective (O_2_) (UPLXAPO40X, Olympus) and a set of relay lenses, and this combination formed a 1.05× imaging system, which approximately met perfect imaging conditions. The remote-focusing module consisted of O_2_ and a 60×1.0NA solid index objective (custom, Special Optics) placed at a 30° angle. The detector arm used dichroic mirrors to simultaneously image all three fluorescence channels. Wide-field imaging used the microscope’s original wide-field optical path.

The illumination light path of the single objective light sheet was also coupled into the microscope. We used a 4-color laser (Colbolt, 405/488/561/633 nm) for illumination and introduced a double-ring modulation (custom chrome plate) on the Fourier plane to generate the Bessel light sheet.

We carried a sample motor stage (U-780, Physik Instrumente) and a live cell workstation (STXG-WSKMX3WX-SET, TokaiHit) on the microscope rack.

### Smart imaging algorithms

The algorithms were composed of two parts: target detection and optomechanical control. The target detection algorithm is mainly developed based on MATLAB. The algorithm uses a three-channel wide-field image as the input and outputs the relative localization of 2025 microchambers and the coordinates of the recommended chambers. To achieve this, first, the brightfield image undergoes two rounds of segmentation using the CellPose network^20^. We set the cell radius to 15 pixels in the first round and 38 in the second round; these correspond to the cell and microchamber radii, respectively. This procedure directly calls the pretrained model and achieves good results without fine-tuning the network. Then, we utilize a secondary template-matching algorithm to localize each chamber. Based on this localization result, the cell segmentation result at the corresponding position is detected, and the cell type and number are determined based on the signal from the fluorescence channel. With a 1-to-1 ratio, all eligible chambers are sorted according to the brightness of the CAR-T cells to provide the recommended results.

The imaging control algorithm is developed based on LabVIEW, which reads the output of the target detection algorithm from the cache file and automatically sets the imaging parameters according to the preset co-positioning relationship between 10× and 60×.

### Super-resolution Algorithm

Our deep learning super-resolution algorithm, developed based on the csbDeep toolkit^33^, was tested on a workstation with an NVIDIA GeForce RTX 4090. A single volume (512×512×380pixels×3chanles) needed approximately 2 minutes to process.

We introduced a deconvolution algorithm for building the training data to improve the network performance. The specific approach was to perform Richardson-Lucy deconvolution on the original data (denoted as I_o_) to obtain the I_d_ based on the measured PSF; then, we refer to ^33^ to train a self-supervised deep learning model to perform isotropic super-resolution on the I_d_ to obtain I_iso_. Finally, using the pair of I_o_ and I_iso_ as the training data, a new network was trained for the super-resolution processing of the data.

### Analyzing Algorithms

To evaluate the reliability of the algorithm, we designed the following experiment: the same batch of CAR-T cells was divided into two groups: one group was treated with the inhibitor dasatinib to simulate impaired cell function, and the other group was left untreated as a control group. Our algorithm performs image analysis of both the experimental group and control group. We compared the algorithm’s results to existing research hypotheses and manual evaluations of the experimental data to test its ability to distinguish normal CAR-T cells from functionally inhibited CAR-T cells.

We developed algorithms based on MATLAB and the packaged front-end as accessible MATLAB apps. We ran the algorithms on a workstation with two CPUs (Intel Xeon Processor E52699 v4 55M Cache 2.20 GHz) and one GPU (NVIDIA GeForce RTX 4090). As an example, scanning 40 chambers produces raw data with a size of 1.2 TB, and 8 hours was needed to visualize and analyze the data.

The algorithms consist of three parts. The first is the reconstruction of the original light-sheet microscopy data. The process can abstract as an affine transformation of the 3D volume. This process took approximately 3 hours.

The second part is to apply a 3D-U-Net to segment the cell structure. The network uses the code from ^34^ with some modifications. The source code is in Python, but we coded a MATLAB interface. We retrained the model, and the 3D manual annotation tool used was ImageJ-Labkit^35^. The training process needed approximately 12 hours, and the prediction of a set of experimental data needed approximately 5 hours.

The third part uses morphology, centroid tracking, and other computer vision algorithms to extract and present image information. This part of the algorithm uses parallelization acceleration and runs on the CPU. Approximately 4 hours was needed to run the first part, but a negligible amount of time was needed to run the second part in parallel with the first part.

