## Supplementary Information for "3D live imaging and phenotyping of the subcellular cytotoxicity in cancer immunotherapy using event-triggered Bessel oblique plane microscopy"

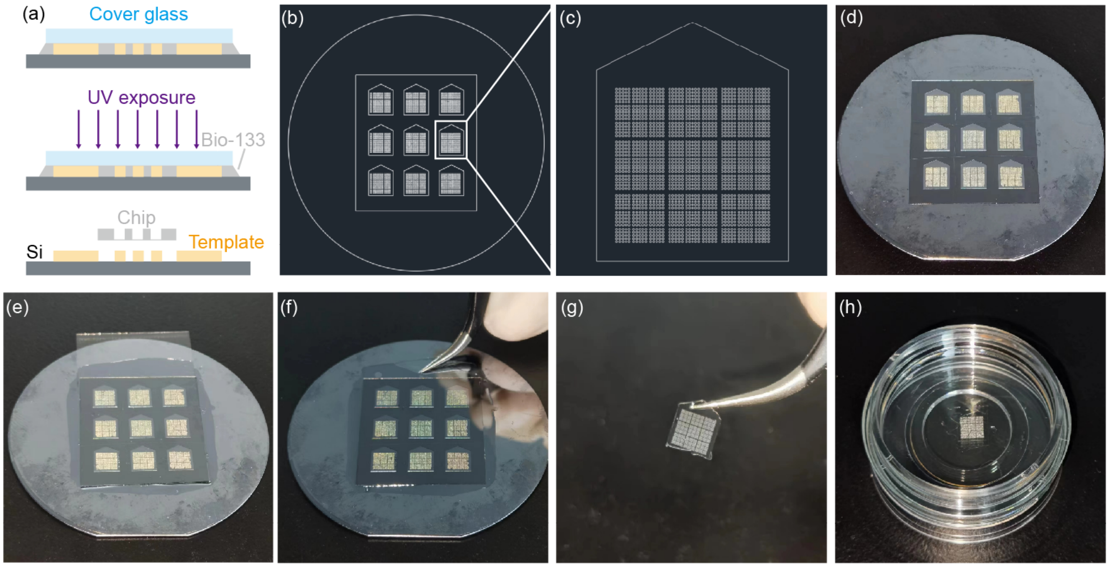


**Supplementary Figure 1. Preparation of the cell-interaction microchip.** (a) Schematic of the Bio-133 chip fabrication process. A layer of uncured Bio-133 (gray) was deposited on a Si wafer containing template structures (yellow), and the wafer was covered with a cover glass. After UV exposure, the cured Bio-133 chip was demolded. (b) Overall chip mold design diagram. (c) Magnified view of the chip design. (d) Template structures fabricated using photoresist SU8-2005. (e) Bio-133 chip fabrication before UV exposure. (f) Demolding of the cured Bio-133 chip. (g) Fabricated Bio-133 chip containing thousands of microwells for hosting the cell pairs. (h) Loading the chip onto a confocal dish.

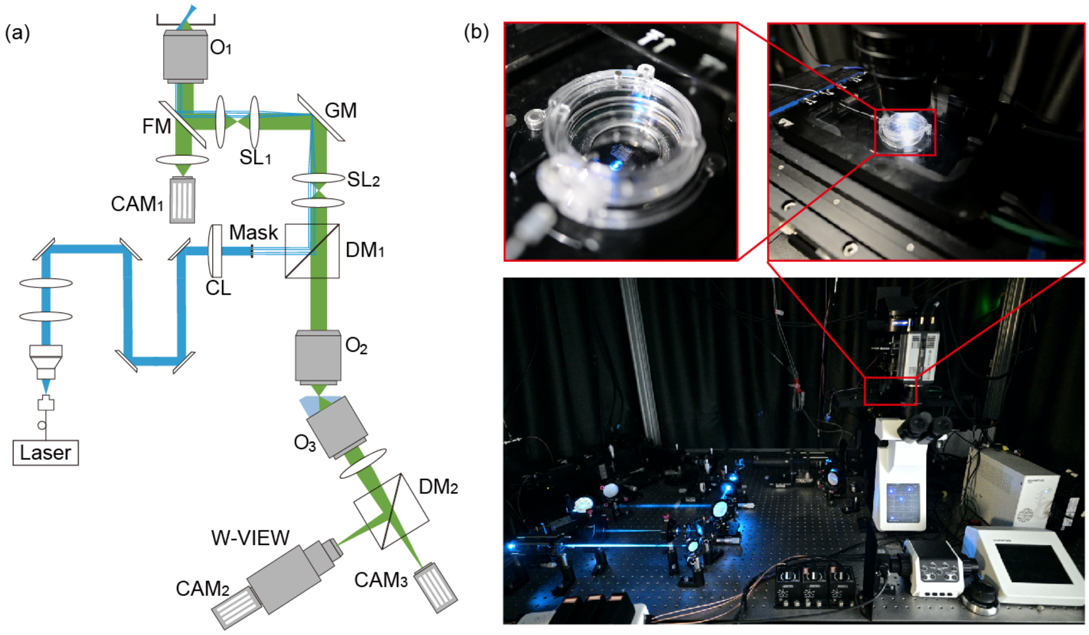


**Supplementary Figure 2. Oblique plane light-sheet microscope (OPM) setup.** (a) Schematic of the laboratory-built OPM built based on a commercial inverted fluorescence microscope (Olympus IX83). Our system contains the following: 1. a single illumination/detection objective convenient for chip-based large-scale imaging and 2. a double-ring modulated Bessel laser sheet with an extended illumination range suitable for imaging float cells. $O_1$: 60×1.3NA silicone oil objective (UPLSAPO60XS2, Olympus). FM: flip mirror inside an Olympus IX83 rack for switching between wide-field and light-sheet imaging. SL: scan lens. GM: galvo mirror. DM: dichroic mirror. CL: cylindrical lens. Mask: A double-ring mask was used to generate a Bessel light sheet[1]. $O_2$: 40×0.95NA air objective (UPLXAPO40X, Olympus). O3: 60×1.0NA solid index objective (custom, Special Optics). $CAM_1$: wide-field camera. $CAM_2$: light-sheet camera. $CAM_3$: light-sheet camera with an image splitter (W-VIEW GEMINI Image Splitting Optics, A12801-01, Hamamatsu). (b) Photographs of the constructed Bessel-OPM system in the working state.

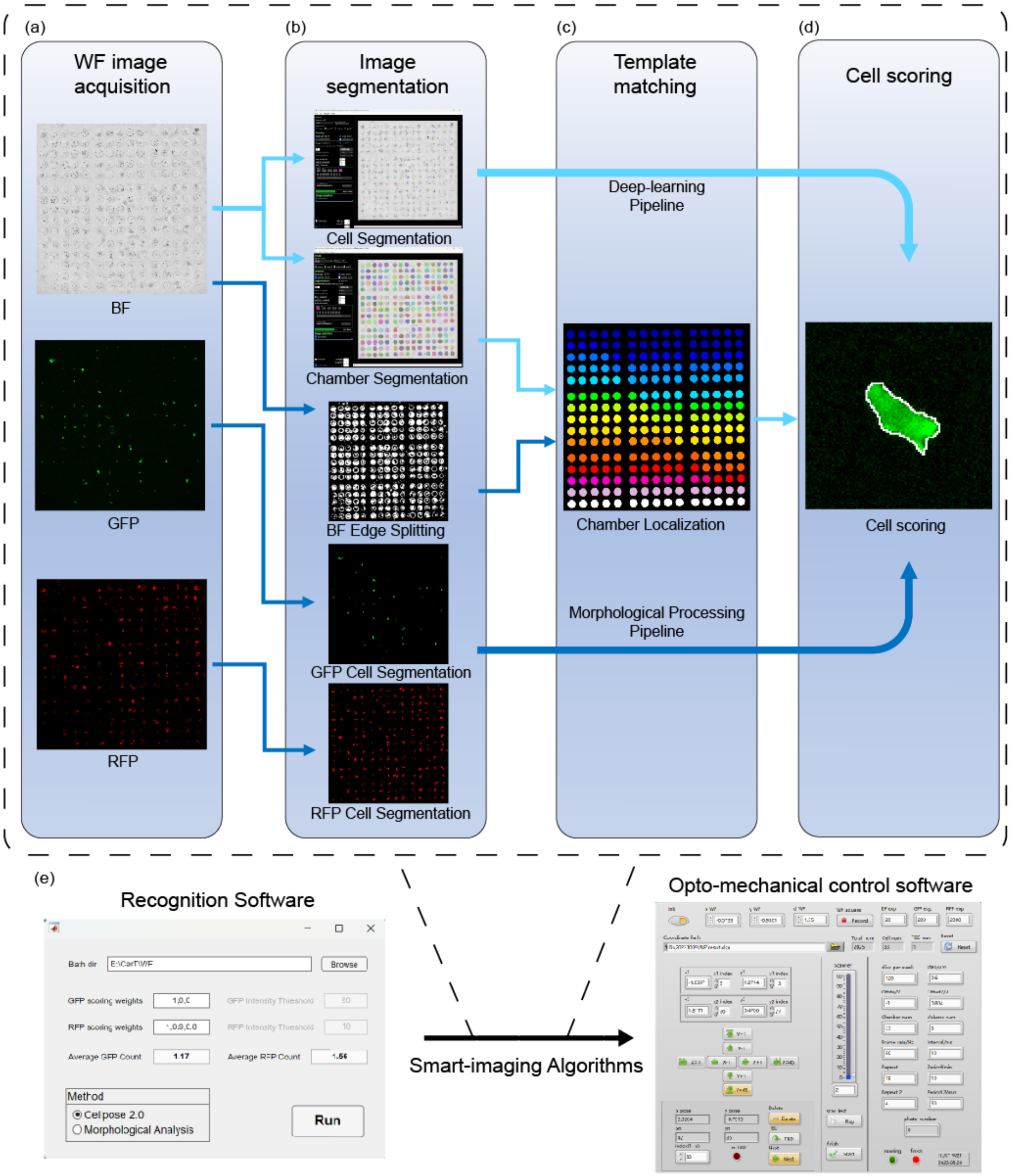


**Supplementary Figure 3. Smart imaging algorithms.** (a) Control wide-field acquisition based on prior chip structure information. (b) Segmentation of the cell mask from the acquired wide-field image. Our algorithm relied on CellPose[2] (cyan arrow in the figure), but an alternative option was to skip the deep learning algorithms and use the morphological algorithms to segment cells (blue arrow in the figure), which was simpler for buildup but had a decrease in accuracy. (c) Identification and localization of chambers by using the template matching algorithm. (d) Cell segmentation and scoring according to the cell number, average fluorescence intensity, and other metrics. (e) Selection of the chambers with the high score and the feedback to the imaging control software for imaging.

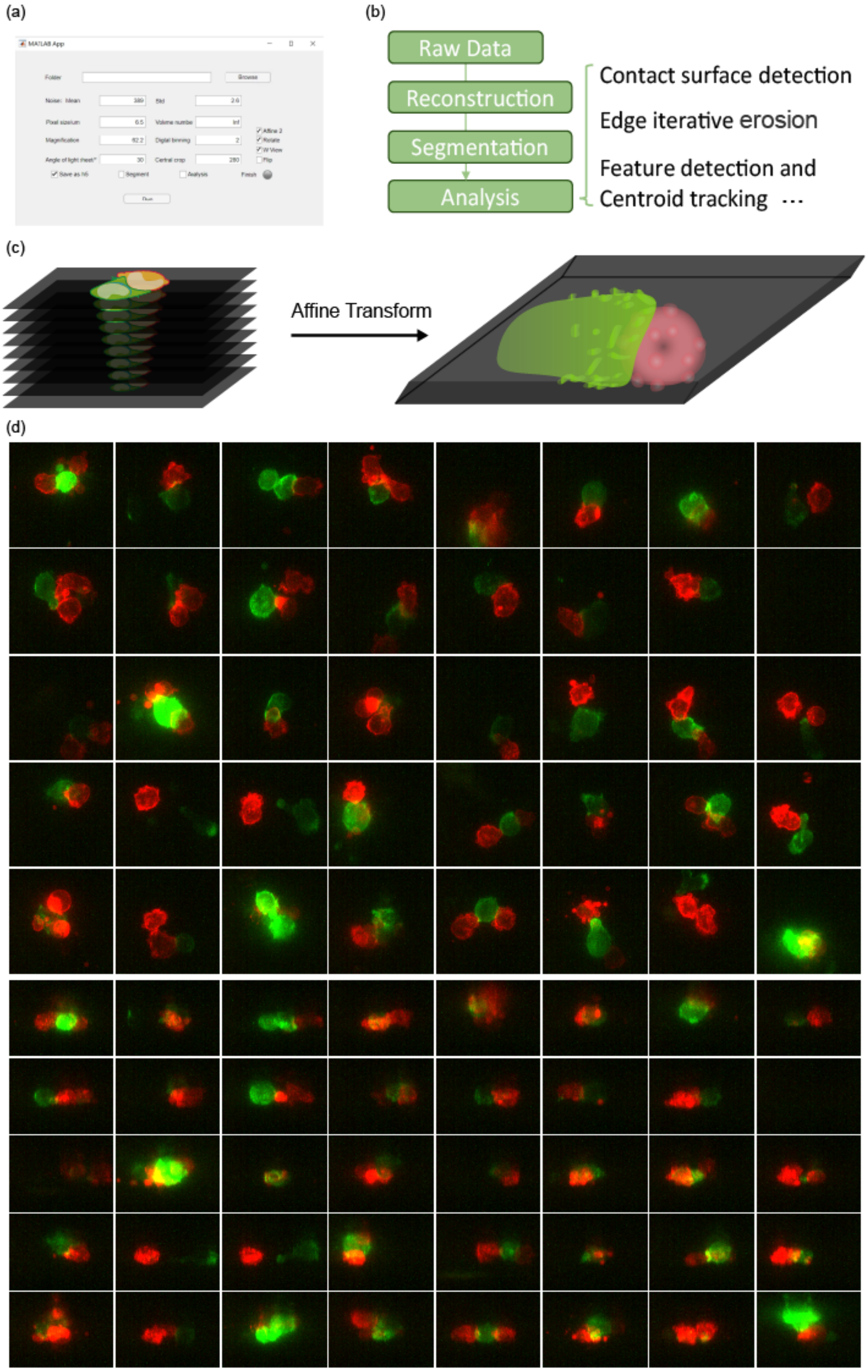


**Supplementary Figure 4. Reconstruction and data analysis pipeline.** (a) Software interface. (b) Overview of the data analysis pipeline. (c) Raw data captured by OPM that were spatially tilted and needed to be reconstructed to their original spatial relationships by three-dimensional affine transformations. (d) First function of our software; batch reconstruct the raw light sheet data and generation of a stitched-together video of the projection map.

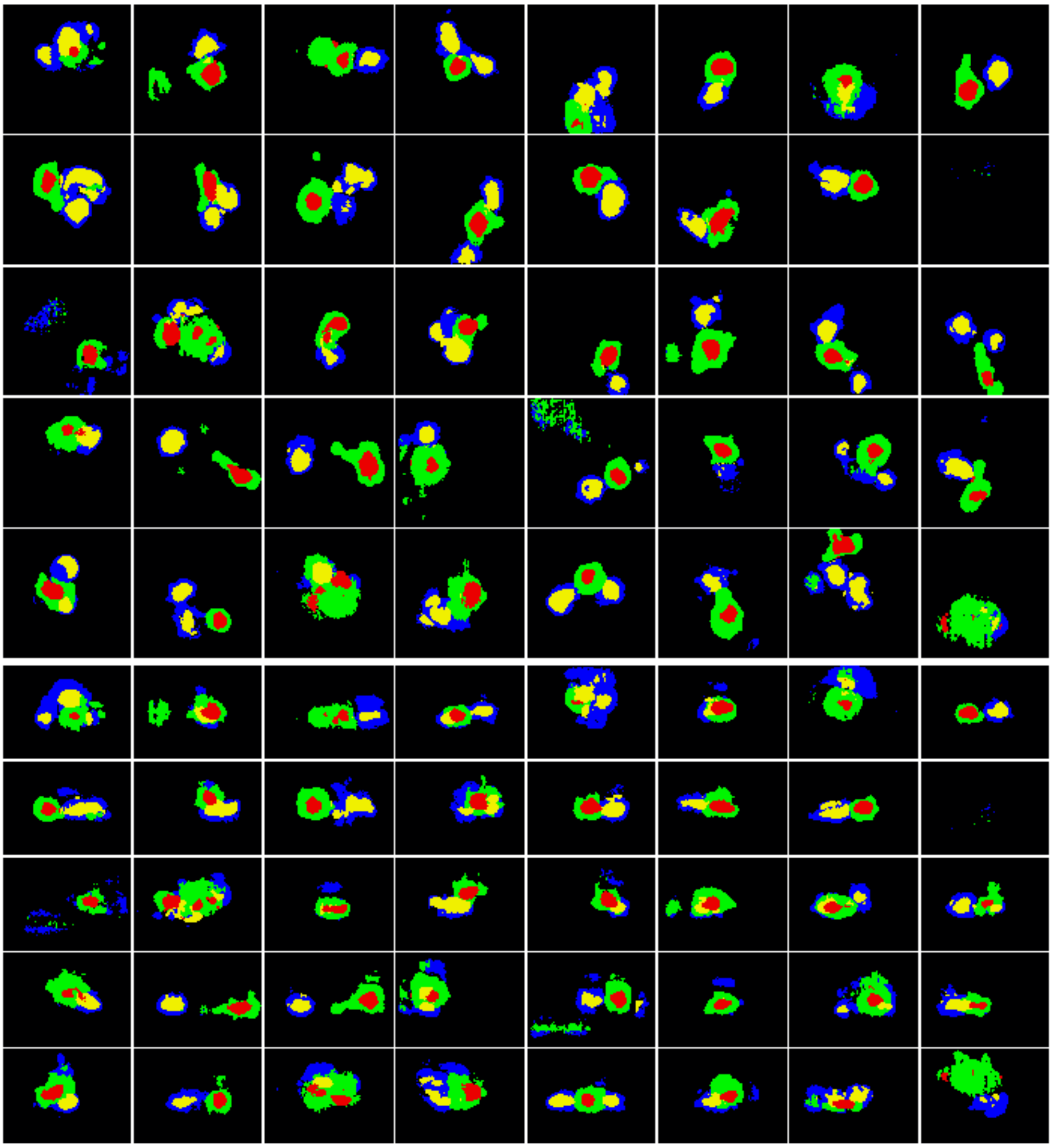

**Supplementary Figure 5. Cell segmentation based on deep learning.** This figure shows the segmentation results corresponding to Supplementary Figure 6(d). In this figure, the green labels indicate CAR-T-cell actin, the blue labels indicate the Nalm6 cell membrane, the red labels indicate the CAR-T-cell nucleus, and the yellow labels indicate the Nalam6 cell nucleus.

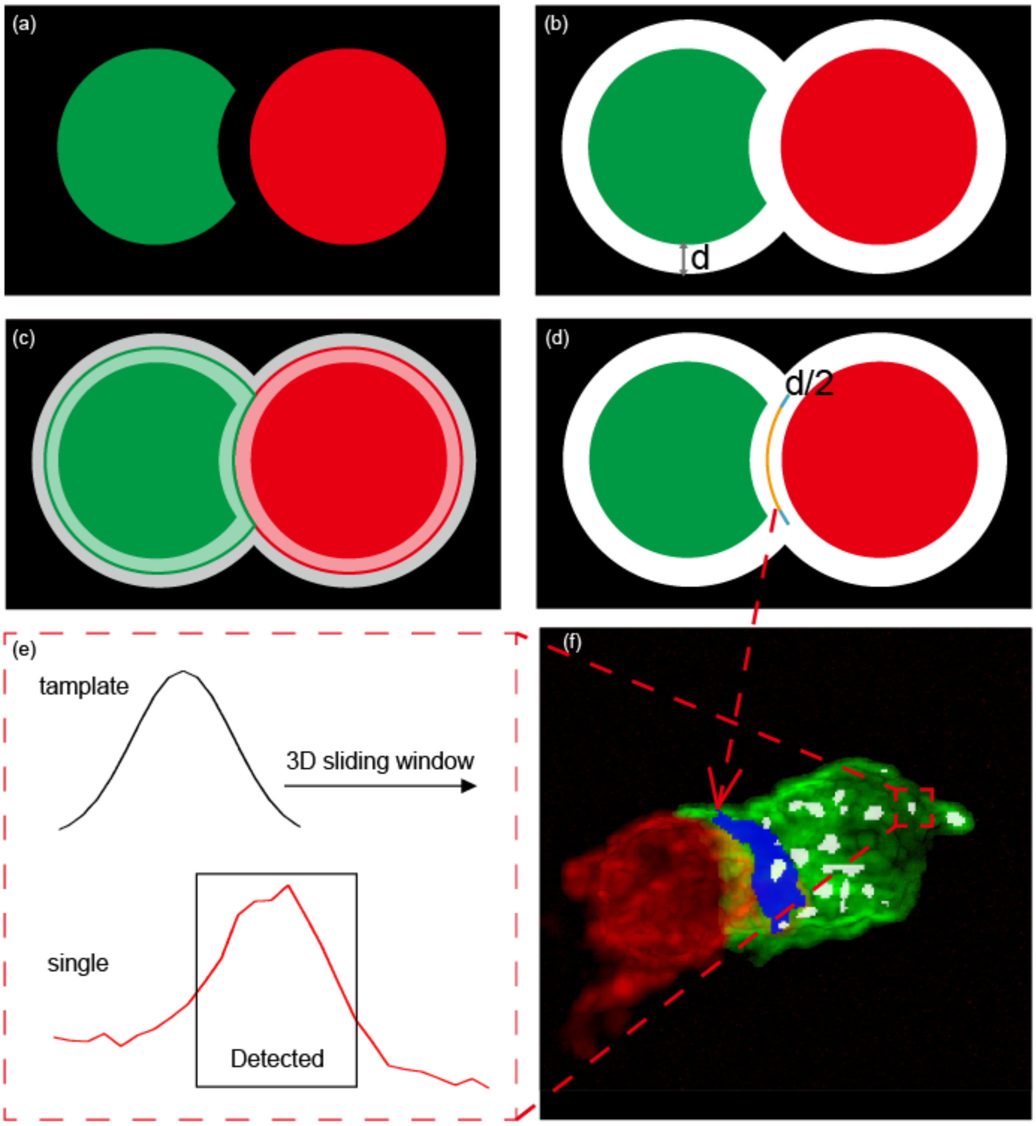


**Supplementary Figure 6. Cytotoxicity analysis algorithms (CAAs) .** (a) ~ (d) Schematic of the contact surface detection algorithm (CSSA). (a) The segmentation results are divided into two categories: one is the mask of CAR-T cells, and the other is the mask of Nalm6 cells; both are eroded by one voxel. (b) Both types of masks are dilated by a distance d, using the union set as the region of interest and the rest as the background. (c) Using the watershed algorithm to segment the intersection lines between three regions (CAR-T, Nalm6 and background regions), only the intersection lines between the CAR-T and Nalm6 regions are used as contact surfaces. (d) The results are cut by d/2 near the background area. By iteratively etching the contact surface with the cutoff part as the seed, we can obtain the distance between each position on the contact surface and the edge of the contact surface. (e) We use the Gaussian correlation detection algorithm to identify the actin feature points. A 3D Gaussian kernel is used to slide on the image, the correlation coefficient r is calculated for each window, and the area with r>0.6 is defined as the feature area. The migration rate of actin is determined based on the centroid tracking results of the feature points. (f) Algorithm segmentation results. Purple represents the contact surface mask, and white represents the feature point mask.

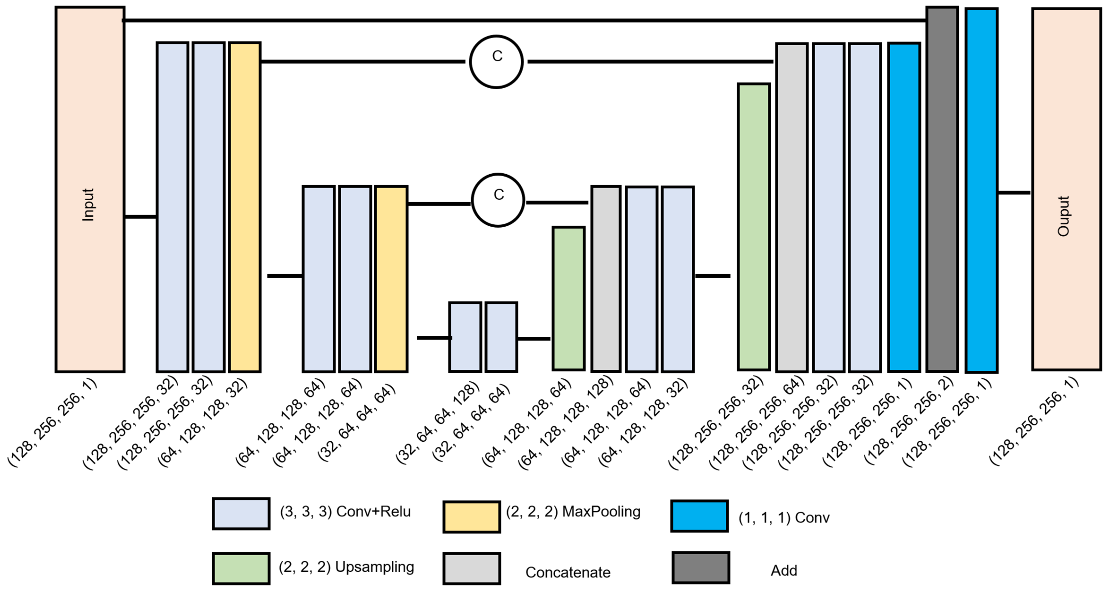


**Supplementary Figure 7. Super-resolution network.** The convolutional neural network utilized in this study follows the architecture of U-Net, consisting of multiple convolutional layers, max-pooling layers, and upsampling layers. The convolutional layers are designed for feature extraction, while the max pooling layers aggregate the spatial information of the features and expand the channel dimension information; in this design, spatial dimensions are reduced by half, and the channel dimensions are doubled. The upsampling layers aim to upsample the downsampled features back to the original feature size, by doubling spatial dimensions while halving channel dimensions. The upsampling method employed here is transposed convolution. Additionally, Concentrate and Add layers are incorporated into the network to reutilize the shallow information, enhancing the network performance. The notation (128, 256, 256, 1) displayed in the figure represents the sizes of four dimensions: z, y, x, and channel, respectively. The sizes of z, y, and x are heuristic values provided for illustrative purposes; in practical network inference, three-dimensional images are automatically segmented into appropriate sizes for inference. After the inference, the inferred small blocks are concatenated to form the final result.

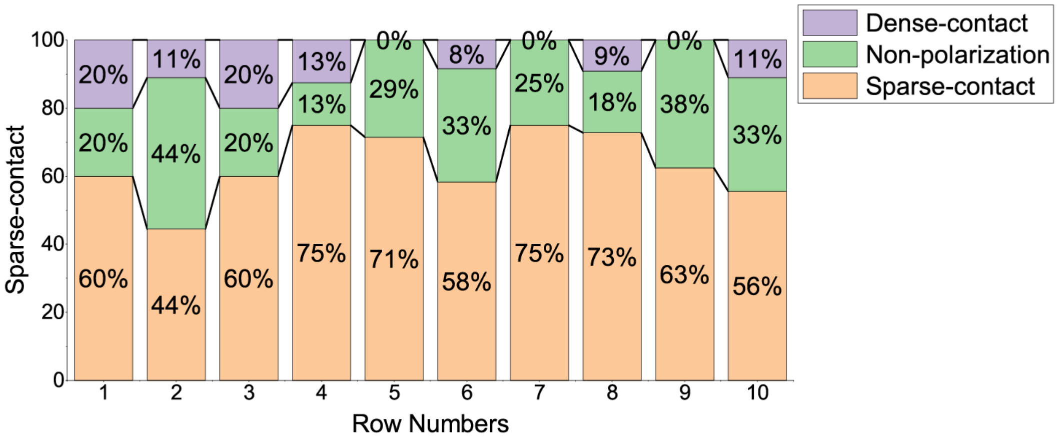


**Supplementary Figure 8. Statistics of the immune synaptic formatting process.** We divided the morphology of CAR-T cells before they contacted Nalm6 cells into two categories. One type was "non-polarization," which does not show significant polarization or form immune synapses. The other type was "polarization," with one end protruding with sparse actin and the other expanding with dense actin. Here, the immune synapses that formed on the sparse actin side were in sparse contact, and those that formed immune synapses on the dense actin side were in dense contact. The percentages of the three cases are shown in the figure. In summary, we captured 78 sets of data on the formation process of immune synapses, with a positive rate of 62.8% and a negative rate of 9.0%.

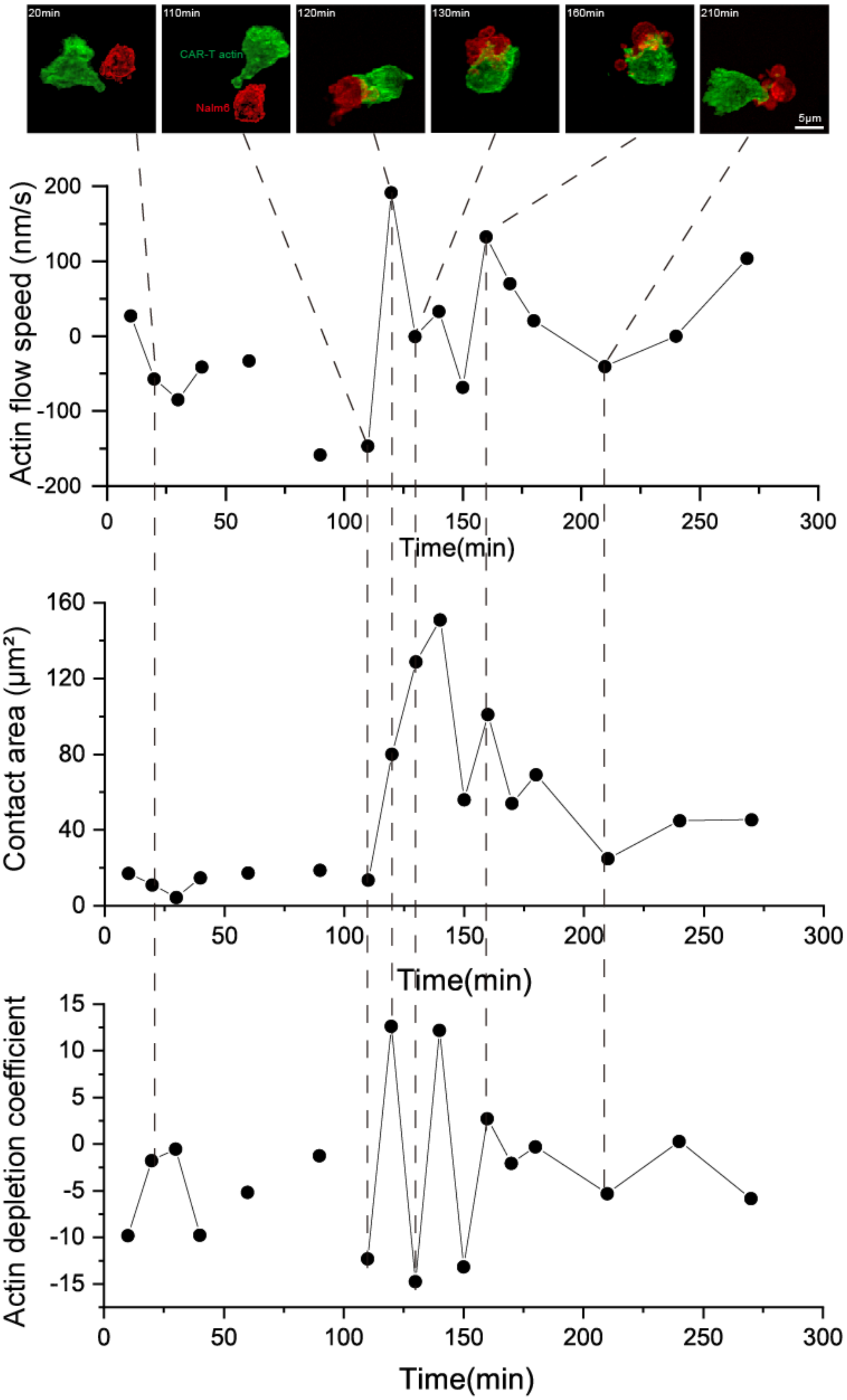


**Supplementary Figure 9. Change in the statistical features in the temporal dimension.** Due to the complexity of the temporal data structure, we did not present the statistical parameters in the main text at each time point. However, in the data with significant changes in CAR-T-cell status, we observed a correlation between the statistical data and immune synaptic formation events.

.

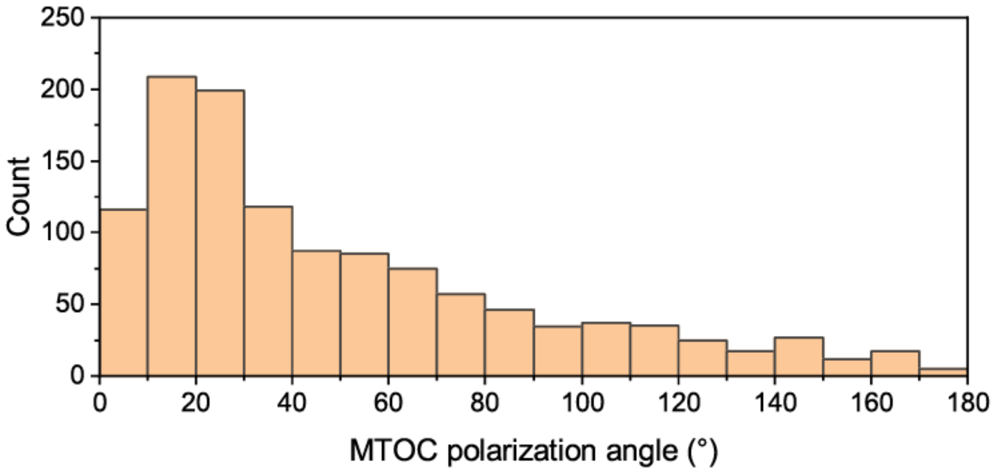


**Supplementary Figure 10. MTOC deviation angle histogram.** In contrast to Fig. 5 (f) in the main text, this figure does not use the minimum value as a representation but instead includes data from all time points. A polarization angle of the centrosome below 90° is still a common trend, indicating that our conclusion is not a deviation generated by the minimum value statistics.

### Supplementary Notes

#### 1. **CAAs Design**

Microscopy is one of the most important research tools in the biomedical field, and its research ability is an important direction for future technological development[3]. The ultimate concept proposed by researchers is to combine microscopy with large deep learning models such that users can automatically complete imaging, analysis and statistical analyses by providing instructions; an example is the automatic imaging and analysis of the area of immune synapses. However, at this stage, the large model technique consumes many arithmetic resources and causing difficulty to perform extensive research[4], while the small model faces many challenges in terms of generalizability and reliability[4]. Therefore, we use small-model deep learning algorithms combined with traditional algorithms to construct intelligent microscopy imaging algorithms that can work robustly in specific scenarios at a lower cost.

Data analysis started with the segmentation of the fluorescence images. In our study, due to the high dynamic range of the fluorescence intensity, traditional adaptive thresholding segmentation algorithms or region-growing-based segmentation algorithms were not able to robustly segment the cells, and manual intervention was needed. In recent years, deep learning segmentation algorithms have been shown to improve these problems. Convolutional neural networks (CNNs) have been proven to have better performance and strong reliability in the field of image segmentation[5], and the U-Net structure is able to effectively converge to avoid overfitting with a small number of training samples[6]. Therefore, in the existing work[7], we used the 3D U-Net network to segment the fluorescence images of cells.

#### 2. **Deep-Learning Segmentation**

The input of the network was a three-channel fluorescence image, and the output was a four-channel segmentation result, which included the actin region of the CAR-T cells, the inner region of the CAR-T cells, the cell membrane of the target cells, and the inner region of the target cells (**Supplementary Fig. 5**). Notably, the image segmentation problem can be subdivided into semantic segmentation problems and instance segmentation problems, and the U-Net structure is only applicable to semantic segmentation problems. The U-Net could only identify whether a voxel belonged to a CAR-T-cell or a target cell but could not distinguish between two different CAR-T cells. By four-channel segmentation, we could perform a watershed segmentation of the image with the inner region of cells as the kernel to achieve instance segmentation. As the smart imaging algorithm selected only microchambers containing one CAR-T-cell with one target cell, the sample segmentation was not considered except for the CAR-T-cell division.

For the training data, we used the ImageJ plugin Labkit[8] for the labeling of 3D annotations. We selected 10 sets of data for annotation; these included high and low signal-to-noise ratios, single multiple cells, irregularly shaped cells, etc., and 8 sets were used as the training dataset and two sets were used as the validation dataset. For the network structure, we used a 4-level U-Net structure with 32, 64, 128, and 256 feature maps. The network converged in approximately 1 hour of training on a single RTX4090 GPU. For high data throughput, we optimized the read/write logic of the original network to increase its capability for parallel reading of a large number of images, and the output weights of the network were not stored but directly exported the binarized segmentation results; this process saved time for subsequent data processing.

#### 3. **Composite Segmentation Algorithm**

In addition to cell segmentation, we used U-Net to complete all image segmentation tasks at once, including immune synaptic (IS) segmentation, microtubule-organizing center (MTOC) segmentation, and dead cell segmentation; however, the output results from the fusion network were not satisfactory. In particular, the network output was ambiguous, and it was difficult for IS segmentation to achieve accurate 3D contour segmentation. Therefore, we separated some simpler or special needs segmentation tasks, and they were used with traditional algorithms.

Segmentation of the IS could not be directly obtained by dilating the segmentation of target cells (SOT) by one voxel and then using the intersection with the segmentation of CAR-T cells (SOC); these cells were not usually closely adjacent to each other due to the ambiguity of the deep learning algorithms. To segment the IS, we designed the segmentation algorithm shown in **Supplementary Fig. 6a~d**.

1. In the first step, we eroded the SOT and SOC by 1 voxel each to ensure that they did not have any direct neighbors. (**Supplementary Fig. 6a**)
2. In the second step, we dilated the eroded SOT and SOC by ***d*** voxels, where ***d*** was the only parameter that needed to be adjusted in this algorithm. This parameter determined that the SOT and SOC was considered in direct contact only when their spacing was less than ***2d***; however, this parameter had very little effect on the algorithm, and not all cells with a spacing of less than ***2d*** were considered in contact. This problem would be compensated for in subsequent algorithmic processes. (**Supplementary Fig. 6b**)
3. In the third step, the intersection of the segmentation of the dilated SOT and SOC was used, and then the complement of the intersection was used to obtain the segmentation of the background region (BR). The pre-segmentation of the BR helped to reduce the amount of computation in the next step of the distance transformation. (**Supplementary Fig. 6c**)
4. In the fourth step, the eroded SOT and SOC in the first step, as well as the BR obtained in the third step, were used as a merged set, and we performed a distance transformation to classify each voxel to the segmentation with its nearest distance; here, the voxels in the entire image volume were classified as one of the three regions: the CAR-T-cell, target cell, and background regions. (**Supplementary Fig. 6c**)
5. In the fifth step, the CAR-T-cell region was dilated by 1 voxel and then intersected with the target cell region. Since the two regions were directly adjacent and did not overlap, the segmentation was a 3D surface with a thickness of 1 voxel. (**Supplementary Fig. 6d**)
6. In the sixth step, since the SOT and SOC were expanded, the IS area derived in the fifth step was large. To solve this problem, we dilated the BR by ***d/2*** voxels and used the intersection with the segmentation result in the fifth step to obtain the IS edge; the remainder of the portion that did not intersect with the background region was the segmentation of the IS. (**Supplementary Fig. 6d**)

Through the algorithm design, we were able to compensate for the maximum ***2d*** voxel error generated by the deep learning segmentation, and only ***d/2*** recompensation was subsequently needed to maximize the robustness and accuracy of the algorithm.

For MTOC segmentation, we used the concept of target detection. According to the average morphological features of the MTOC, we defined a Gaussian kernel of 11*11*11 voxels with a standard deviation of 3.5 voxels. By traversing the 3D image volume using the Gaussian kernel sliding window, each time the window slid, the Pearson correlation coefficient between the values of voxels in the window and the Gaussian kernel was computed, and if the correlation coefficient was greater than +0.6, then it was assumed that this point might be part of the MTOC (**Supplementary Fig. 6e**). Since the correlation operation was time-consuming, we additionally specified an intensity threshold for MTOC, and voxels with fluorescence signals below the threshold were skipped for correlation detection, which greatly improved the speed of the operation.

Cell death was determined by the fluorescence intensity of SYTOX™ Blue dye. Since the signal from fluorescence microscopy was quantitative, we used a fixed threshold segmentation for dead cell segmentation; specifically, the cell was considered dead when the fluorescence signal of the blue fluorescent channel inside the cell was greater than the threshold value.

#### 4. Quantitative Algorithm

Based on the above segmentation results, we designed subsequent algorithms for analyzing the morphological and behavioral characteristics of the cells. Although the performance of deep learning algorithms is progressing, for specific physiological phenomena, the analysis results based entirely on deep learning algorithms are still not universally recognized at this stage due to their black-box nature; however, traditional algorithms have the advantages of clear logic and well-documented intermediate results in this regard. Therefore, we did not introduce deep learning methods in the quantification and analysis of cellular features.

##### 4.1 Computation of the area of the IS

Since we ensure a single voxel-thickness segmentation of the IS, the number of voxels of the IS segmentation was counted, and these could be converted to the area of the IS. Notably, the same voxel corresponds to an actual area ranging from 1 unit to $\sqrt{2}$ units due to the difference in the spatial angle of the 3D surface at that voxel. Since the order-of-magnitude did not differ, to improve the algorithm's running efficiency, 1 voxel was regarded as a 1-unit contact area for all algorithms used in this study.

##### 4.2 Determination of the IS center and the coefficient between the IS center distance and actin fluorescence intensity

The algorithm utilized IS edge segmentation in the fifth step of the composite segmentation algorithm.

1. In the first step, the IS edge was used as the ***seed***.
2. In the second step, the ***seed*** was dilated by 1 voxel, and the intersection of the IS segmentation was used to obtain the secondary edge segmentation (SES).
3. In the third step, the voxel of the SES was assigned the value of ***i*** and saved to the resultant volume, where ***i*** was the number of iterations.
4. In the fourth step, the seed was replaced by the SES,
5. In the fifth step, the SES was subtracted from the IS segmentation, and then it was determined whether the IS segmentation was empty; if it was empty, then the iteration ended; otherwise return to the second step.

After the above algorithm, we obtained the distance from each voxel on the IS to the edge of the IS, and the voxel farthest from the edge was defined as the center of the IS. Subsequently, the distance of these voxels was collected with the actin fluorescence intensity information, and Pearson's correlation coefficient was calculated.

##### 4.3 Reflux velocity of actin

In this study, we observed that actin reflux was not linear and that actin tended to reflux but was also accompanied by random oscillations in the vertical reflux direction; therefore, we chose to quantify the actin reflux velocity by calculating the projected velocity using three-dimensional vector analysis.

1. In the first step, we defined the axis of interaction (AOI); this was the vector from the center of the IS, pointing toward the center of mass of the segmentation of the CAR-T-cell (COC).
2. In the second step, a Gaussian kernel was defined, and the correlation coefficient was calculated using an algorithm similar to that for segmenting the MTOC, thus segmenting the dendritic structure of the actin. (**Supplementary Fig. 6e**)
3. In the third step, we traced the structures.
4. In the fourth step, the traces with larger displacements were screened, and the projections of their displacement vectors on the AOI were averaged.

The traces with larger displacements were screened because a large number of structures at the edge of the IS were easily recognized by the algorithm as dendritic structures; moreover, if they were not screened, the statistical results would be invalid. Notably, the cell itself drifted within the imaging time window; here, we incorporated a compensation mechanism into the algorithm that tracked the displacement of the COC, and this displacement was subtracted from all actin displacement vectors to ensure that the vectors only reflected the motion of the actin itself relative to the cell.

##### 4.4 MTOC polarization angle

We defined the vector pointing from the MTOC to the COC as the axis of the MTOC (AOM), and the angle between the AOM and the AOI was the MTOC polarization angle, which ranged from 0 to 180°. Notably, we used the spatial angle to measure the MTOC polarization based on the a priori fact that the center of CAR-T cells is occupied by the nucleus and that the MTOC usually does not pass through the nucleus but moves at the edge of the cell; therefore, the polar coordinates can intuitively and clearly describe the MTOC polarization.

#### Supplementary Videos

**Supplementary Video 1. The working pipeline of EBOPM and CAAs.**

**Supplementary Video 2. The trans-scale smart imaging strategy in EBOPM.**

**Supplementary Video 3. The 5D visualization of CAR-T cytotoxic processes obtained by the high-throughput mode of EBOPM.**

**Supplementary Video 4. The 5D visualization of CAR-T cytotoxic processes suppressed by Dasatinib treatment.**

**Supplementary Video 5. The Establishment process of immune synapses captured by the continuous mode of EBOPM.**
